# Quantitative systems pharmacological analysis of drugs of abuse reveals the pleiotropy of their targets and the effector role of mTORC1

**DOI:** 10.1101/470922

**Authors:** Fen Pei, Hongchun Li, Bing Liu, Ivet Bahar

**Affiliations:** Department of Computational and Systems Biology, School of Medicine, University of Pittsburgh, PA, 15213, USA

**Keywords:** Drug abuse, quantitative systems pharmacology, pleiotropic proteins, mTOR complex 1, drug-target interactions, neurotransmission, machine learning, cellular pathways, addiction development

## Abstract

Existing treatments against drug addiction are often ineffective due to the complexity of the networks of protein-drug and protein-protein interactions (PPIs) that mediate the development of drug addiction and related neurobiological disorders. There is an urgent need for understanding the molecular mechanisms that underlie drug addiction toward designing novel preventive or therapeutic strategies. The rapidly accumulating data on addictive drugs and their targets as well as advances in machine learning methods and computing technology now present an opportunity to systematically mine existing data and draw inferences on potential new strategies. To this aim, we carried out a comprehensive analysis of cellular pathways implicated in a diverse set of 50 drugs of abuse using quantitative systems pharmacology methods. The analysis of the drug/ligand-target interactions compiled in DrugBank and STITCH databases revealed 142 known and 48 newly predicted targets, which have been further analyzed to identify the KEGG pathways enriched at different stages of drug addiction cycle, as well as those implicated in cell signaling and regulation events associated with drug abuse. Apart from synaptic neurotransmission pathways detected as a common upstream signaling module that ‘senses’ the early effects of drugs of abuse, pathways involved in neuroplasticity are distinguished as determinants of neuronal morphological changes. Notably, many signaling pathways converge on important targets such as mTORC1. The latter is proposed to act as a universal effector of the persistent restructuring of neurons in response to continued use of drugs of abuse.

## 1 Introduction

Drug addiction is a chronic relapsing disorder characterized by compulsive, excessive, and self-damaging use of drugs of abuse. It is a debilitating condition that potentially leads to serious physiological injury, mental disorder and death, resulting in major health and social economic impacts worldwide (Nestler, 2013; Koob and Volkow, 2016). Substances with diverse chemical structures and mechanisms of action are known to cause addiction. Except for alcohol and tobacco, substances of abuse are commonly classified into six groups based on their primary targets or effects: cannabinoids (e.g. cannabis), opioids (e.g. morphine, heroin, fentanyl), central nervous system (CNS) depressants (e.g. pentobarbital, diazepam), CNS stimulants (e.g. cocaine, amphetamine), hallucinogens (e.g. ketamine, lysergic acid diethylamide) and anabolic steroids (e.g. nandrolone, oxymetholone).

The primary actions of drugs of abuse have been well studied. In spite of the pleiotropy and heterogeneity of drugs of abuse, they share similar phenotypes: from acute intoxication to chronic dependence (Taylor et al., 2013), the reinforcement shift from positive to negative through a three-stage cycle involving binge/intoxication, withdrawal/negative effect, and preoccupation/anticipation (Koob and Volkow, 2016). Notably, virtually all drugs of abuse augment dopaminergic transmission in the reward system (Wise, 1996). However, the detailed cellular pathways of addiction processes are still far from known. For example, cocaine acts primarily as an inhibitor of dopamine (DA) transporter (DAT) and results in DA accumulation in the synapses of DA neurons (Shimada et al., 1991; Volkow et al., 1997). However, it has been shown that DA accumulation *per se* is not sufficient to account for the rewarding process associated with cocaine addiction; serotonin (5-HT) and noradrenaline/norepinephrine (NE) also play important roles (Rocha et al., 1998; Sora et al., 1998). Another example is ketamine, a nonselective antagonist for N-methyl-d-aspartate (NMDA) receptor (NMDAR), notably most effective in the amygdala and hippocampal regions of neurons (Collingridge et al., 1983). In addition to its primary action, ketamine affects a number of other neurotransmitter receptors, including sigma-1 (Mendelsohn et al., 1985), substance P (Okamoto et al., 2003), opioid (Hustveit et al., 1995), muscarinic acetylcholine (mACh) (Hirota et al., 2002), nicotinic acetylcholine (nACh) (Coates and Flood, 2001), serotonin (Kapur and Seeman, 2002), and γ-aminobutyric acid (GABA) receptors (Hevers et al., 2008). The promiscuity of drugs of abuse brings an additional layer of complexity, which prevents the development of efficient treatment against drug addiction.

In recent years there has been significant progress in the characterization of drug/target/pathway relations driven by the accumulation of drug-target interactions and pathways data, as well as the development of machine learning, *in silico* genomics, chemogenomics and quantitative systems pharmacology (QSP) tools. Several innovative studies started to provide valuable information on substance abuse targets and pathways. For example, Li et al. curated 396 drug abuse related genes from the literature and identified five common pathways underlying the reward and addiction actions of cocaine, alcohol, opioids and nicotine (Li et al., 2008). Hu et al. analyzed the genes related to nicotine addiction via a pathway and network-based approach (Hu et al., 2018). Biernacka et al. performed genome-wide analysis on 1165 alcohol-dependence cases and identified two pathways associated with alcohol dependence (Biernacka et al., 2013). Xie et al. generated chemogenomics knowledgebases focused on G-protein coupled receptors (GPCRs) related to drugs of abuse in general (Xie et al., 2014), and cannabinoids in particular (Xie et al., 2016). Notably, these studies have shed light on selected categories or subgroups of drugs. There is a need to understand the intricate couplings between multiple pathways implicated in the cellular response to drugs of abuse, identify mechanisms common to various categories of drugs while distinguishing those unique to selected categories.

We undertake here such a systems-level approach using a dataset composed of six different categories of drugs of abuse. Following a QSP approach proposed earlier (Stern et al., 2016), we provide a comprehensive, unbiased glimpse of the complex mechanisms implicated in addiction. Specifically, a set of 50 drugs of abuse with a diversity in chemical structures and pharmacological actions were collected as probes, and the known targets of these drugs as well as the targets predicted using our probabilistic matrix factorization (PMF) method (Cobanoglu et al., 2013) were analyzed to infer biological pathways associated with drug addiction. Our analysis yielded 142 known and 48 predicted targets and 173 pathways permitting us to identify both generic mechanisms regulating the responses to drug abuse as well as specific mechanisms associated with selected categories, which could both facilitate the development of auxiliary agents for treatment of addiction.

A key step in our approach is to identify the targets for drugs of abuse. There exists various drug-target interaction databases (DBs), web servers and computational models, as summarized recently (Chen et al., 2016). The drug-target interaction DBs utilized in this work are DrugBank (Wishart et al., 2018) and STITCH (Szklarczyk et al., 2016). DrugBank is a bioinformatics and cheminformatics resource that combines drug data with comprehensive target information. It is frequently updated, with the current version containing 10,562 drugs, 4,493 targets and corresponding 16,959 interactions. Since most of drugs of abuse are approved or withdrawn drugs, DrugBank is a good source for obtaining information on their interactions. STITCH, on the other hand, is much more extensive. It integrates chemical-protein interactions from experiments, other DBs, literature and predictions, resulting in data on 430,000 chemicals and 9,643,763 proteins across 2,031 genomes. We have used the subset of human protein-chemicals data supported by experimental evidence. The method of approach adopted here is an important advance over our original PMF-based machine learning methodology for predicting drug-target interactions (Cobanoglu et al., 2013). First, the approach originally developed for mining DrugBank has been extended to analyzing the STITCH DB, the content of which is 2-3 orders of magnitude larger than DrugBank (based on the respective numbers of interactions). Second, the information on predicted drug-target associations is complemented by pathway data on *humans* inferred from the KEGG pathway DB (December 2017 version) (Kanehisa et al., 2017) upon pathway enrichment analysis of known and predicted targets. Third, the outputs are subjected to extensive analyses to detect recurrent patterns and formulate new hypotheses for preventive or therapeutic strategies against drug abuse.

## 2 Materials and Methods

### 2.1 Selection of drugs of abuse and their known targets

We selected as input 50 drugs commonly known as drugs of abuse using two basic criteria: (i) diversity in terms of structure and mode of action, and (ii) availability of information on at least one human target protein in DrugBank v5 (Wishart et al., 2018) or STITCH v5 (Szklarczyk et al., 2016). The selected drugs represent six different categories: CNS stimulants, CNS depressants, opioids, cannabinoids, anabolic steroids and hallucinogens (see **Supplementary Table 1** and **Supplementary Figure 1** for details).

A dataset of 142 known targets, listed in **Supplementary Table 2**, were retrieved from DrugBank and STITCH DBs for these 50 drugs. The list includes all targets reported for these drugs in DrugBank, and those with high confidence score, based on experiments, reported in STITCH. Each chemical-target interaction is annotated with five confidence scores in STITCH: experimental, DB, text-mining, prediction, and a combination score of the previous four, each ranging from 0 to 1. We selected the human protein targets with experimental confidence scores of 0.4 or higher. **Supplementary Table 2** summarizes the 142 targets we identified as well as the associated 445 drug-target interactions.

Structure-based and interaction-pattern-based similarities between pairs of drugs were evaluated using two different criteria. The former was based on *structure-based distance* calculated as the Tanimoto distance between their 2D structure fingerprints. The Tanimoto distances were evaluated using Python RDKit suite (RDKit: Open-Source Cheminformatics Software. https://www.rdkit.org/). Similarities based on their interactions patterns with known targets were evaluated by evaluating *target-based distances*. To this aim, we represented each drug *i* by a 142-dimensional ‘target vector’ ***d**_i_*, the entries of which represent the known targets and are assigned values of 0 or 1, depending on the existence/observation of an interaction between the corresponding target and drug *i*. Interaction-pattern similarities between drug pairs *i* and *j* were evaluated by calculating the correlation cosine cos(***d_i_*** · ***d_j_***) = (***d_i_***. ***d_j_***) / (|***d_i_***| |***d_j_***|) between these vectors, and the corresponding cosine distance is [1 - cos(***d_i_*** · ***d_j_***)]. Likewise, *ligand-based distances* between target pairs *i* and *j* were evaluated as the cosine distance between the 50-dimensional vectors ***t_i_*** and ***t_j_*** corresponding to the two targets, the entries of which are 0 or 1 depending on absence or existence of an interaction between the target and the corresponding drug of abuse.

### 2.2 Probabilistic matrix factorization (PMF) based drug-target interaction prediction

Novel targets for each drug were predicted using our probabilistic matrix factorization (PMF) based machine learning approach (Cobanoglu et al., 2013; Cobanoglu et al., 2015). Briefly, we start with a sparse matrix ***R*** representing the known interactions between *N* drugs and *M* targets. Using the PMF algorithm, we decomposed ***R*** into a drug matrix ***U*** and a target matrix ***V***, by learning the optimal *D* latent variables to represent each drug and each target. The product of ***U^T^*** and ***V*** assigns values to the unknown (experimentally not characterized) entries of the reconstructed ***R***, each value representing the *confidence score* for a novel drug-target interaction prediction

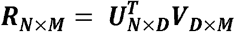

Using this method, we trained two PMF models, one based on 11,681 drug-target interactions between 6,640 drugs and 2,255 targets from DrugBank v5, and the other based on 8,579,843 chemical-target interactions for 311,507 chemicals and 9,457 targets from STITCH v5 human experimentally confirmed subset, respectively. We evaluated the confidence scores in the range [0, 1] for each predicted drug-target interaction, in both cases. We selected the interactions with confidence scores higher than 0.7 within the top 10 predicted targets for each input drug. This led to 161 novel interactions identified between 27 out of the 50 input drugs and 89 targets (composed of 41 known and 48 novel targets) (**Supplementary Table 3**).

### 2.3 Pathway enrichment analysis

We mapped the 50 drugs with 142 known and 48 predicted targets to the KEGG pathways (version December 2017, *homo sapiens*) (Kanehisa et al., 2017). 114 and 173 pathways were mapped by 142 known targets and all targets (both known and predicted) respectively (see **Supplementary Table 4**). In order to prioritize enriched pathways, we calculated the hypergeometric *p*-values based on the targets as the enrichment score as follows. Given a list of targets, the enrichment *p*-value for pathway *A* (*P^A^*) is the probability of randomly drawing *k_0_* or more targets that belong to pathway *A:*

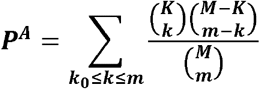

where *M* is the total number of human proteins in the KEGG Pathway, *m* is the total number of proteins/targets we identified, and *K* is the number of proteins that belong to pathway *A*, while *k_0_* is the number of targets we identified that belong to pathway A.

## 3 Results

### 3.1 Functional similarity of drugs of abuse does not imply structural similarity, consistent with the multiplicity of their actions

**Figure 1** presents a quantitative analysis of the functional and structural diversity of the examined *n* = 50 drugs of abuse, as well as the similarities of their *m* = 142 targets. The *n* × *n* maps in panels **A** and **B** display the drug-drug pairwise distances/dissimilarities based on their 2D fingerprints (panel **A**), and their interaction patterns with their targets (panel **B**). Panels **C** and **D** display the corresponding dendrograms. The drugs are indexed and color-coded as in **Supplementary Table 1** and **Supplementary Figure 1**. As expected, drugs belonging to the same functional category (same color) exhibit more similar interaction patterns (panel **D**). However, we also note outliers, such as cocaine lying among opioids, as opposed to its categorization as a CNS stimulant, or promethazine, a CNS depressant, lying among hallucinogens (shown by *arrows*). The peculiar behavior of cocaine is consistent with its high promiscuity (see **Figure 2A** for the number of targets associated with each examined drug). This type of promiscuity becomes even more apparent when the drugs are organized based on their structure (or 2D fingerprints; see *Materials and Methods*) as may be seen in panel **A.** For example, opioids (clustered together in panels **B** and **D** based on their interactions) are now distributed in two or more branches of the dendrogram (*cyan labels/arc*; panel **C**); likewise, CNS depressants (*blue*) and cannabinoids (*light brown*), grouped each as a single cluster in panel **D**, are now distributed into two or more clusters in panel **C**.

**Figure 1.**
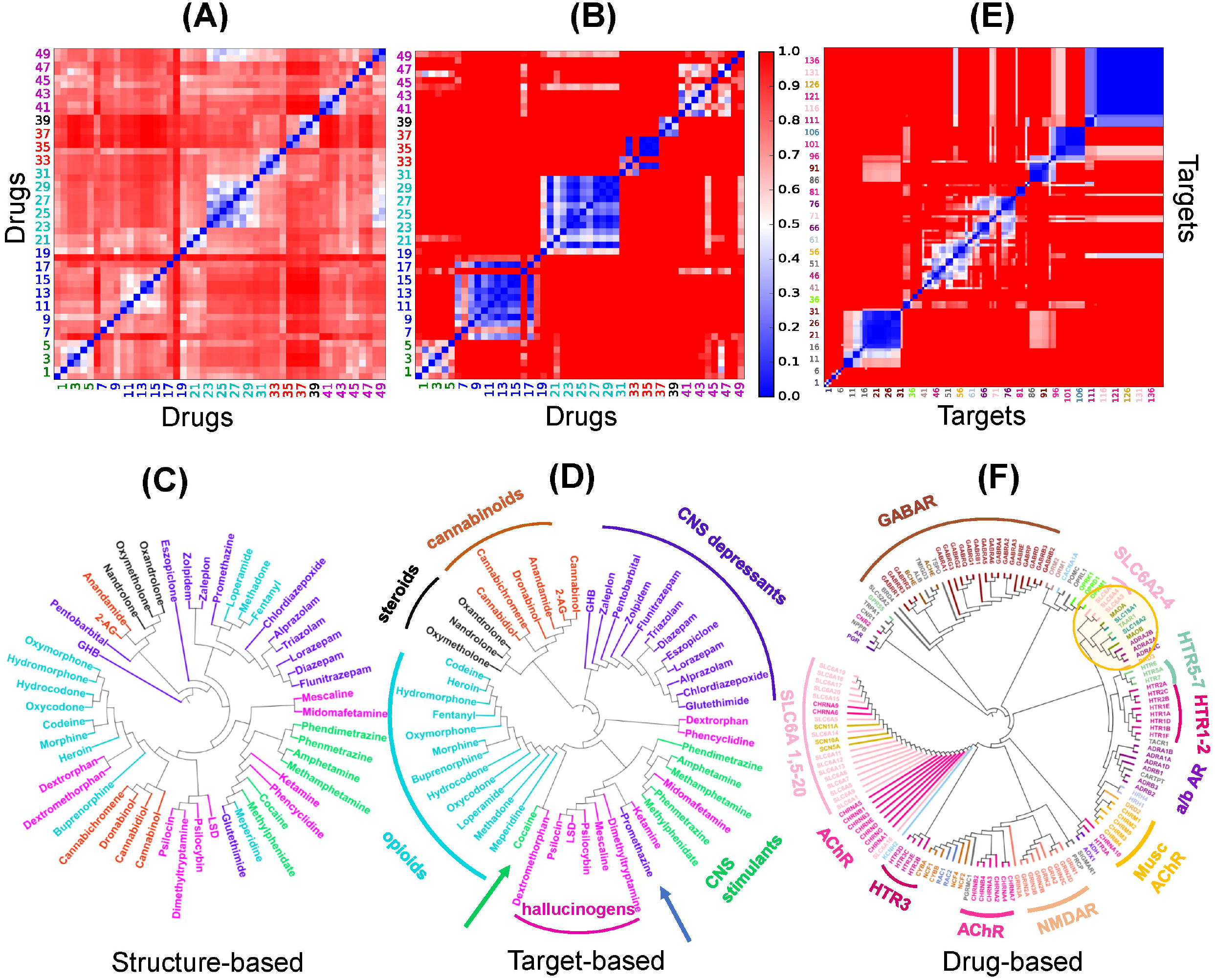
Distribution of the dataset of 50 drugs of abuse based on their structure and interaction (with targets) similarities (A-D), and pairwise similarities and classification of the corresponding targets based on their interaction patterns with the drugs of abuse. **(A-D)** Drug-drug distance maps for the studied 50 addictive drugs based on **(A)** 2D structure fingerprints and **(B)** interaction patterns with targets using the correlation cosines between their target vectors (see *Materials and Methods*), and corresponding dendrograms **(C-D)**. The indices of drugs of abuse in **(A)** and **(B)** follow the same order as those used in **Supplementary Table 1**. The drug labels in **C** and **D** are color-coded based on their categories: CNS stimulants (green), CNS depressants (blue), opioids (cyan), cannabinoids (*light brown*), anabolic steroids (*black*) and hallucinogens (magenta). Note that the drugs of abuse in the same category do not necessarily show structural similarities nor similar interaction pattern with targets. **(E)** Pairwise distance map for the 142 known targets based on their interaction patterns with the 50 drugs. The indices in **(E)** follows the same order as those listed clockwise in the dendrogram **(F)**. The tree maps in **(C)**, **(D)** and **(F)** are generated based on the respective distances values in the **(A)**, **(B)** and **(E)**.

**Figure 2.**
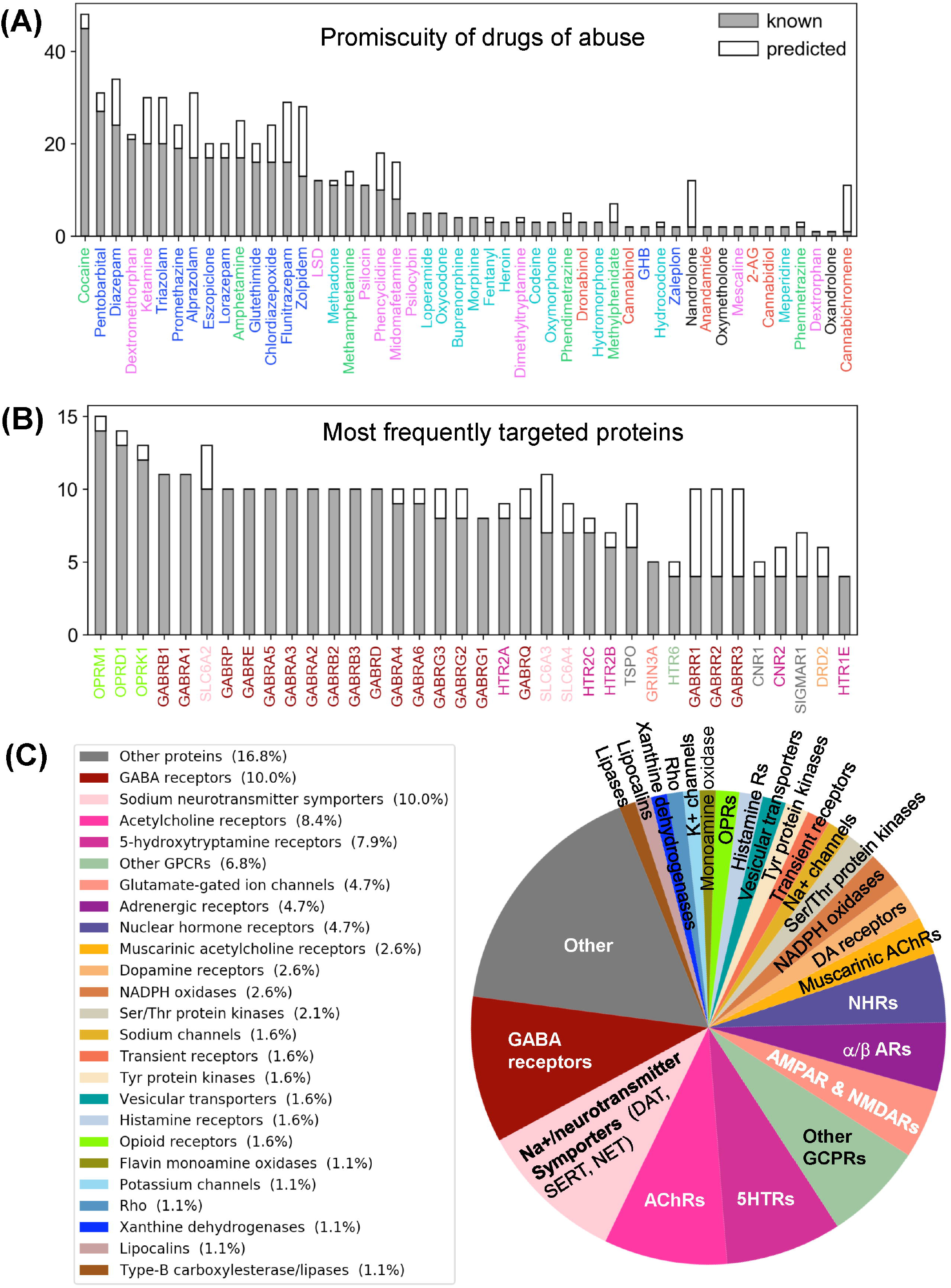
Promiscuity of drugs of abuse and their targets, and major families of proteins targeted by drugs of abuse. Number of known (*gray*) and predicted (*white*) interactions are shown by bars for **(A)** drugs of abuse and **(B)** their targets. The examined set consists of 50 drugs of abuse and a total of 142 known and 48 predicted targets, involved in 445 (known) and 161 (predicted) interactions. Panel **A** displays the number of interactions known or predicted for all 50 drugs. Panel **B** displays the results for the targets that interact with at least 4 known drugs (36 targets). The colors used for names of drugs and targets are same as those used in **Figure 1**. Panel C displays the distribution of families of proteins targeted by drugs of abuse.

Overall these results suggest that the functional categorization of the drugs does not necessarily comply with their structural characteristics. The similar functionality presumably originates from targeting similar pathways, but the difference in the structure suggests that either their targets, or the binding sites on the same target, are different; or the binding is not selective enough such that multiple drugs can bind the same site. Consequently, a diversity of pathways or a multiplicity of cellular responses are triggered by the use and abuse of these drugs.

### 3.2 The selected drugs and identified targets are highly diverse and promiscuous

We evaluated the similarities between proteins targeted by drugs of abuse, based on their interaction patterns with the studied drugs of abuse. **Figure 1** panels **E** and **F** display the respective target-target distances, and corresponding dendrogram. **Supplementary Table 2** lists the full names of these targets, organized in the same order as the panel **E** axes. We discern several groups of targets clustered together in consistency with their biological functions. For example, practically all GABA receptor subtypes (*brown*) are clustered together. This large cluster also includes the riboflavin transporter 2A (SLC52A2), which may be required for GABA release (Tritsch et al., 2012). On the other hand, the different subtypes of serotonin (or 5-hydroxytryptamine, 5-HT) receptors (5HTRs) participate in distinct clusters pointing to the specificity of different subtypes vis-à-vis different drugs of abuse (*labeled* in **Figure 1F**).

The large majority of neurotransmitter transporters, such as Na^+^/Cl^-^-dependent GABA transporters (SLC6A1) and glycine transporter (SLC6A9) are in the same cluster (*pink*, *labeled*). Acetylcholine receptors also lie close to (or are even interspersed among) Na^+^/Cl^-^-dependent neurotransmitter transporters, presumably due to shared drugs such as cocaine. However, the three transporters playing a crucial role in developing drug addiction, DAT, NE transporter (NET) and serotonin transporter (SERT) (*labeled* SLC6A2: NET, SLC6A3: DAT, SLC6A4: SERT) are distinguished by from all other neurotransmitter transporters as a completely disjoint group. The corresponding branch of the dendrogram (*highlighted by the yellow circle*) also includes vesicular amino acid transporters and trace amine-associated receptor 1 (TAAR1) known to interact with these transporters (Miller, 2011). We also note in the same branch two seemingly unrelated targets: flavin monoamine oxidase which draws attention to the role of oxidative events; and α2-adrenergic receptor subtypes A-C, which uses NE as a chemical messenger for mediating stimulant effects such as sensitization and reinstatement of drug seeking, and adenylate cyclase as another messenger to regulate cAMP levels (Sofuoglu and Sewell, 2009).

**Supplementary Table 2** summarizes the 445 known interactions between these 50 drugs and 142 targets. We observe an average of 8.9 interactions per drug and 3.1 interactions per target. There are 23 promiscuous drugs that target at least 10 proteins as shown in **Figure 2** panel **A**. Cocaine, the most promiscuous psychostimulant, interacts with 45 known and 3 predicted targets. It is known that cocaine binds DAT to lock it in the outward-facing state (OFS) and block the reuptake of DA. It similarly antagonizes SERT and NET (Heikkila et al., 1975; Sora et al., 1998), and also affects muscarinic acetylcholine receptors (mAChRs) M1 and M2 (Williams and Adinoff, 2008). Our PMF model also predicted a potential interaction between cocaine and M5. While this interaction is not listed in current DBs, there is experimental evidence suggesting that M5 plays an important role in reinforcing the effects of cocaine (Fink-Jensen et al., 2003), in support of the PMF model prediction.

The PMF model enables us to predict novel targets. For example, anabolic steroid nandrolone has only two known interactions, and cannabinoid cannabichromene has one. However, 10 new targets were predicted with high confidence scores for each of them (**Supplementary Table 3** and **Supplementary Figure 2A**). This is due to the data available in STITCH DB, which offers a large training dataset that enhances the performance of our machine learning approach. Overall, 89 new interactions were predicted for known targets, and 42 novel targets were predicted with 72 interactions. **Figure 2** panel **C** displays the distribution of all targets among different protein families. As will be further elaborated below, among the newly identified drug-target pairs, nandrolone-MAPK14 (mitogen-activated protein kinase 14, also known as p38α) and canabichromene-IKBKB (inhibitor of NFκ-B kinase subunit β) play a role in regulating mTORC1 signaling, which will be shown to be an effector of drug addiction.

Turning to targets, three opioid receptors (OPRM1, OPRD1 and OPRL1) exhibit the highest level of promiscuity (**Supplementary Figure 2B**). The μ-type opioid receptor (OPRM1) interacts with 14 known drugs including all opioids as well as ketamine and dextromethorphan. We also predicted a novel interaction between OPRM1 and the CNS stimulant methylphenidate. This is consistent with experimental observations that methylphenidate upregulates OPRM1’s activity in the reward circuitry in a mouse model (Zhu et al., 2011). Furthermore, tissue-based transcriptome analysis (Uhlén et al., 2015) shows that 69% of our 190 targets are expressed in the brain, and 49 of them show elevated expression levels in the brain compared to other tissue types (**Supplementary Table 5**). Among all the targets, NMDA receptor 1 (GRIN1) shows the highest elevated expression. It is also one of the top 5 enriched genes overall in the brain (Uhlén et al., 2015).

Taken together, the 50 selected drugs of abuse and the 142 known and 48 novel targets we identified cover a diversity of biological functions, are involved in many cellular pathways, and are generally promiscuous. In order to reveal the common mechanisms that underlie the development and escalation of drug addiction and also distinguish the effects specific to selected drugs, we proceed now to a detailed pathway analysis, presented next.

### 3.3 Pathway enrichment analysis reveals the major pathways implicated in various stages of addiction development

Our QSP analysis yielded a total of 173 pathways, including 114 associated with the known targets of the examined dataset of drugs of abuse, and 59 associated with the predicted targets. The detailed pathway enrichment results can be found in **Supplementary Table 4**. These pathways can be grouped in five categories (**Figure 3**, **Supplementary Figure 4**, and **Supplementary Table 4**):

**Figure 3.**
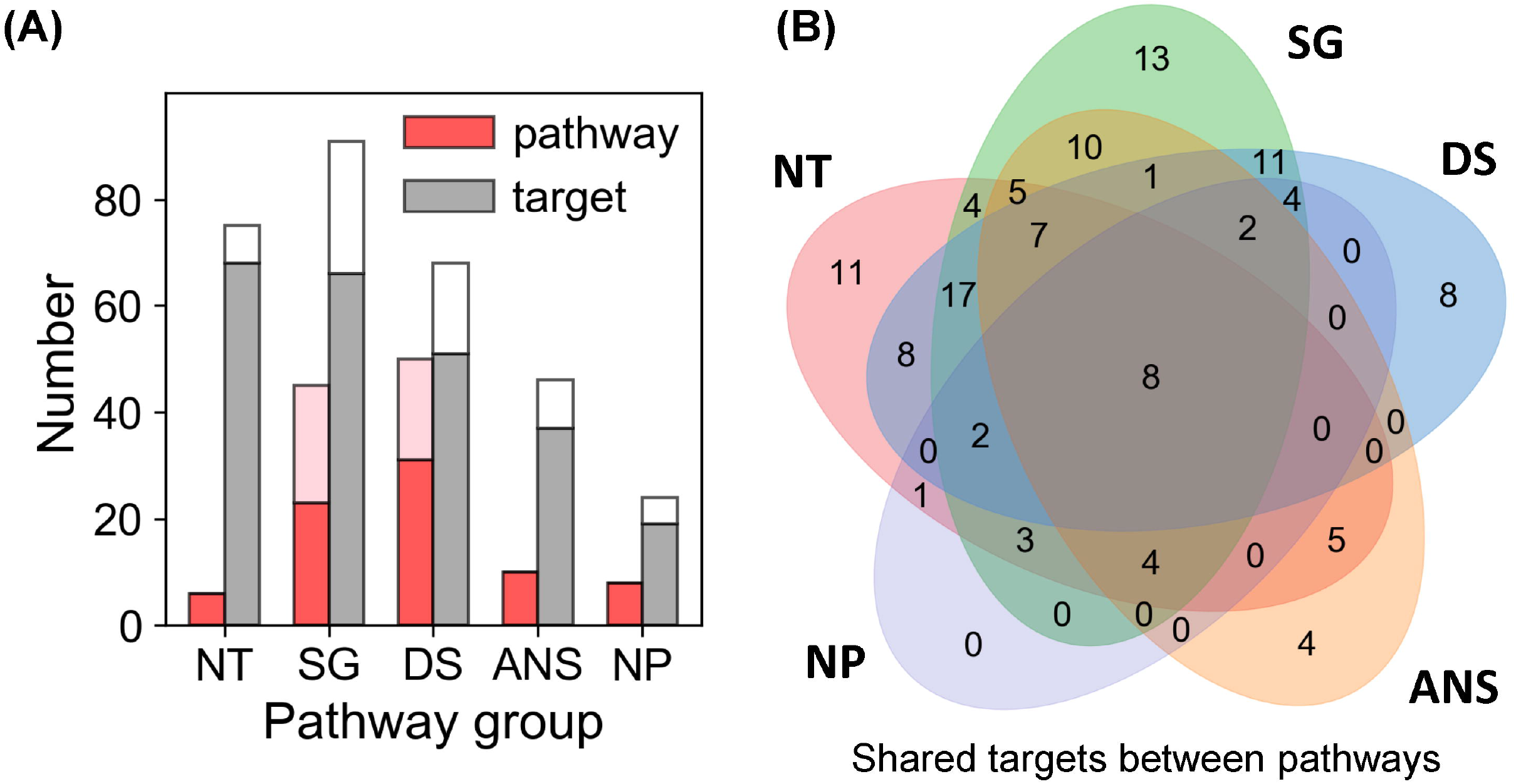
Results from pathway and target enrichments analysis. Five broad categories of pathways are distinguished among those involving the targets of drug abuse: NT, synaptic neurotransmission pathways; SG, signal transduction pathways; DS, disease-associated pathways; ANS, autonomic nervous system-innervation pathways; and NP, neuroplasticity related pathways. **(A)** Numbers of pathways (*red bars*) and targets (*gray bars*) of drug abuse lying in the five categories, based on data available in DrugBank and STITCH. The *pink* and *white* stacked bars are the corresponding numbers for pathways and targets additionally predicted by PMF. **(B)** Overlaps between the target content of the five pathway categories. See the complete list of pathways and targets in **Supplementary Table 4**.

#### Synaptic Neurotransmission (NT)

Six significantly enriched (with *p*-value < 0.05) pathways are associated with synaptic neurotransmission: dopaminergic, serotonergic, glutamatergic, synaptic vesicle cycle, cholinergic, and GABAergic synapses pathways. 68 known targets and 7 predicted targets are involved in these pathways. This is consistent with the fact that neurotransmission plays a dominant role in the rewarding system and is key to drug addiction (Volkow and Morales, 2015).

#### Signal Transduction (SG)

46 intracellular signaling pathways were mapped by 92 targets comprised of 66 known and 25 predicted targets. Notably, many of these pathways have been reported to play a role in mediating the effects of drugs of abuse. These include the top five (calcium signaling (Li et al., 2008), retrograde endocannabinoid signaling (Mechoulam and Parker, 2013), cGMP-PKG signaling (Shen et al., 2016), cAMP signaling (Philibin et al., 2011), and Rap1 signaling (Cahill et al., 2016)) as well as some pathways with relatively low enrichment score (i.e. 0.2 < *p*-value < 0.5), such as TNF signaling (Zhu et al., 2018), MAPK signaling (Sun et al., 2016), PI3K-Akt signaling (Neasta et al., 2011), NF-ĸB signaling (Nennig and Schank, 2017), and mTOR signaling (Neasta et al., 2014). We note that many receptors targeted by drugs of abuse take part in the KEGG neuroactive ligand-receptor interaction pathway. In the interest of focusing on intracellular signaling effects, we have not included these in the SG category; they are listed in the ‘Other Pathways’ in **Supplementary Table 4**.

#### Autonomic nervous system (ANS)-innervation (ANS)

We also identified 10 pathways regulating ANS-innervated systems such as endocrine secretion, taste transduction, and circadian entrainment. Recent evidences suggested drugs of abuse such as morphine (Al-Hasani and Bruchas, 2011) and cocaine (Moeller et al., 1997; Prosser et al., 2014) can influence ANS-innervated systems and may contribute to the withdrawn symptoms associated with drug addiction. 37 known and 9 predicted targets take part in these pathways.

#### Neuroplasticity (NP)

Eight enriched pathways with potential to alter the morphology of neurons, were found to be related to drug addiction. Among them, long-term potentiation (LTP) and long-term depression (LTD) are key to reward-related learning and addiction by modifying the fine tuning of dopaminergic firing (Jones and Bonci, 2005). Axon guidance pathway regulates the growth direction of neuron cells (Bahi and Dreyer, 2005). Regulation of actin cytoskeleton plays important role in morphological development and structural changes of neurons (Luo, 2002). Gap junctions connect neighboring neurons via intercellular channels that allow direct electrical communication (Belousov and Fontes, 2013) and regulate the efficiency of communication between electrical synapses (Belousov and Fontes, 2013). Insulin-like growth factor 1 receptor (IGF1R) is predicted as a target of drug triazolam (**Supplementary Table 4**). IGF1R is involved in LTP, adherens junction and focal adhesion pathways. It functions via canonical signaling pathways noted above in the SG category, such as the PI3K-Akt-mTOR and Ras-Raf-MAPK pathways (Lee et al., 2016) and it plays important role in neuroplasticity (Lee et al., 2016).

#### Disease-associated pathways (DS)

50 enriched pathways are associated with diverse diseases in different organs such as brain, liver, and lung. They also cover various drug addiction mechanisms including: nicotine addiction, morphine addiction, cocaine addiction, amphetamine addiction, and alcoholism. Additionally, there are ‘other pathways’ such as those involved in cell migration, differentiation, immune responses, and metabolic events, which can be seen in **Supplementary Table 4**.

Taken together, the enrichment analysis reveals five major categories of pathways that regulate the three stages of drug addiction cycle: (1) binge and intoxication, (2) withdrawal and negative affect, and (3) preoccupation and anticipation (or craving) (Koob and Volkow, 2010). Drugs of abuse directly affect neurotransmission pathways: they increase the accumulation of DA and other neurotransmitters in the synaptic and extrasynaptic regions, which in turn results in the hedonic feeling (stage 1) and triggers the DA reward system. Dysregulation of ANS-innervation pathways may cause negative effects and feelings (stage 2) and feedback to the CNS. Addictive drugs impair executive processes by disrupting the reward system (neurotransmission pathways) and imparting morphological changes via neuroplasticity pathways (e.g. LTD and LTP), which then result in craving (stage 3). Below, we present an in-depth analysis of the role of these pathways or their shared targets in drug addiction.

### 3.4 Selected targets shared by dominant pathways emerge as common mediators of drug addiction

We next analyzed the overlapping targets between the pathways in different functional categories. We note in particular eight pleiotropic proteins involved in all five categories (at the intersection of the 5 Venn diagrams in **Figure 3B**: AMPA receptor (subtype GluA2; GRIA2), NMDA receptors 1 and 2A-D (designated as GRIN1, GRIN2A, GRIN2B, GRIN2C and GRIN2D) and voltage-dependent calcium channel Ca_v_2.1 (or CACNA1A) as well as the predicted target phosphatidylinositol 3-kinase class 1A catalytic subunit α (PIK3CA) (**Supplementary Table 4**). Additionally, 15 proteins are distinguished as targets of four of these major pathways: Serotonin receptors 5HTR2-A, -B and -C), GABA_A_ receptors 1-6 (GABRA1-GABRA6), β-1 adrenergic receptor 1 (ADRB1), Ras-related C3 botulinum toxin substrate 1 (RAC1; member of Rho family of GTPases), mAChR M_3_ (CHRM3) and DA receptor D_2_ (DRD2) and two predicted targets p38α (MAPK14) and DA receptor D_1_ (DRD1).

AMPA receptor plays a crucial role in LTP and LTD, which are vital to neuroplasticity, memory and learning (Volkow et al., 2016). Serotonin receptors, expressed in both the CNS and the peripheral nervous system (e.g. gastrointestinal tract), are responsible for anxiety, impulsivity, memory, mood, sleep, thermoregulation, blood pressure, gastrointestinal motility and nausea (Pytliak et al., 2011). They have been proposed to be therapeutic targets for treating cocaine use disorder (Howell and Cunningham, 2015). RAC1 is involved in five neuroplasticity pathways, including axon guidance, adherens junction and tight junction pathways (**Supplementary Table 4**), and 13 intracellular signal transduction pathways. It regulates neuroplasticity, as well as apoptosis and autophagy (Natsvlishvili et al., 2015). DA receptor D_2_ is a target of 28 drugs of abuse (out of 50 examined here) and is involved in cAMP signaling, and gap junction pathways, in addition to dopaminergic signaling. It is implicated in reward mechanisms in the brain (Blum et al., 1996) and the regulation of drug-seeking behaviors (Edwards et al., 2006). Finally, PI3K turns out to be the most pleiotropic target among those targeted by drugs of abuse, being involved in 61 pathways identified here, including neuroplasticity pathways such as axon guidance, and several downstream signaling pathways such as PI3K-Akt, mTOR, Ras and Jak-STAT pathways.

Overall, 23 proteins are distinguished here as highly pleiotropic proteins involved in at least four of the five major categories of pathways implicated in drug abuse. Most of them are ligand- or voltage-gated ion channels or neurotransmitter receptors, mainly AMPAR, NMDAR, Cav2.1, mAChR, and serotonin and DA receptors. However, it is interesting to note the targets PI3K and p38α, not currently reported in the DBs DrugBank and STITCH, emerge as highly pleiotropic targets of the drugs of abuse. These are predicted to directly or indirectly affect addiction development. Finally, a number of proteins take part in specific drug-abuse-related pathways and might serve as targets for selective treatments. **Supplementary Table 6** provides a list of such targets uniquely implicated in distinctive pathways.

### 3.5 Pathway enrichment highlights the interference of drugs of abuse with synaptic neurotransmission

It is broadly known that neurotransmitters such as DA, 5-HT, NE, endogenous opioids, ACh, endogenous cannabinoids, Glu and GABA are implicated in drug addiction (Tomkins and Sellers, 2001; Everitt and Robbins, 2005; Parolaro and Rubino, 2008; Benarroch, 2012). Our analysis also showed the serotonergic synapse (*p-value = 4.64E-20*), GABAergic synapse (*p-value = 3.45E-19*), cholinergic synapse (*p-value = 1.64E-08*), dopaminergic synapse (*p-value = 1.25E-07*) and glutamatergic synapse (*p-value = 1.83E-04*) pathways were significantly enriched (**Supplementary Table 4**). A total number of 34 drugs (across six different groups) target at least one of these pathways. However, the identification of a pathway does not necessarily mean that the drug directly affects that particular neurotransmitter transport/signaling. There may be indirect effects due to the crosstalks between synaptic signaling pathways. For example, the ionotropic glutamate receptors NMDAR and AMPAR are also the downstream mediators in the dopaminergic synapse pathway. Likewise, GABARs are downstream mediators in the serotonergic synapse pathway.

In **Figure 4**, we highlight five major neurotransmission events that directly mediate addiction, and illustrate how eight drugs of abuse interfere with them. Despite the promiscuity of the drugs of abuse, some selectively map onto a single synaptic neurotransmission pathway. For example, psilocin (a hallucinogen whose structure is similar to 5HT (Diaz and Diaz, 1997)) interacts with several types of 5HTRs, regulating serotonergic synapse exclusively (see **Figure 4** and **Supplementary Table 4**). In contract, loperamide (not shown) affects all neurotransmission pathways by interacting with the voltage-dependent P/Q-type calcium channel (VGCC), regulating calcium flux on synapses. Cocaine targets four of these synaptic neurotransmission events (serotonergic, GABAergic, cholinergic, and dopaminergic synapses), through its interactions with 5-HT3R, sodium- and chloride-dependent GABA transporter (GAT), muscarinic (M1 and M2) and nicotinic AChRs, and DAT, respectively. Methadone affects three synaptic neurotransmissions, including serotonergic synapse, dopaminergic synapse and glutamatergic synapse through the interactions with SERT, DAT, and glutamate receptors (NMDAR) respectively.

**Figure 4.**
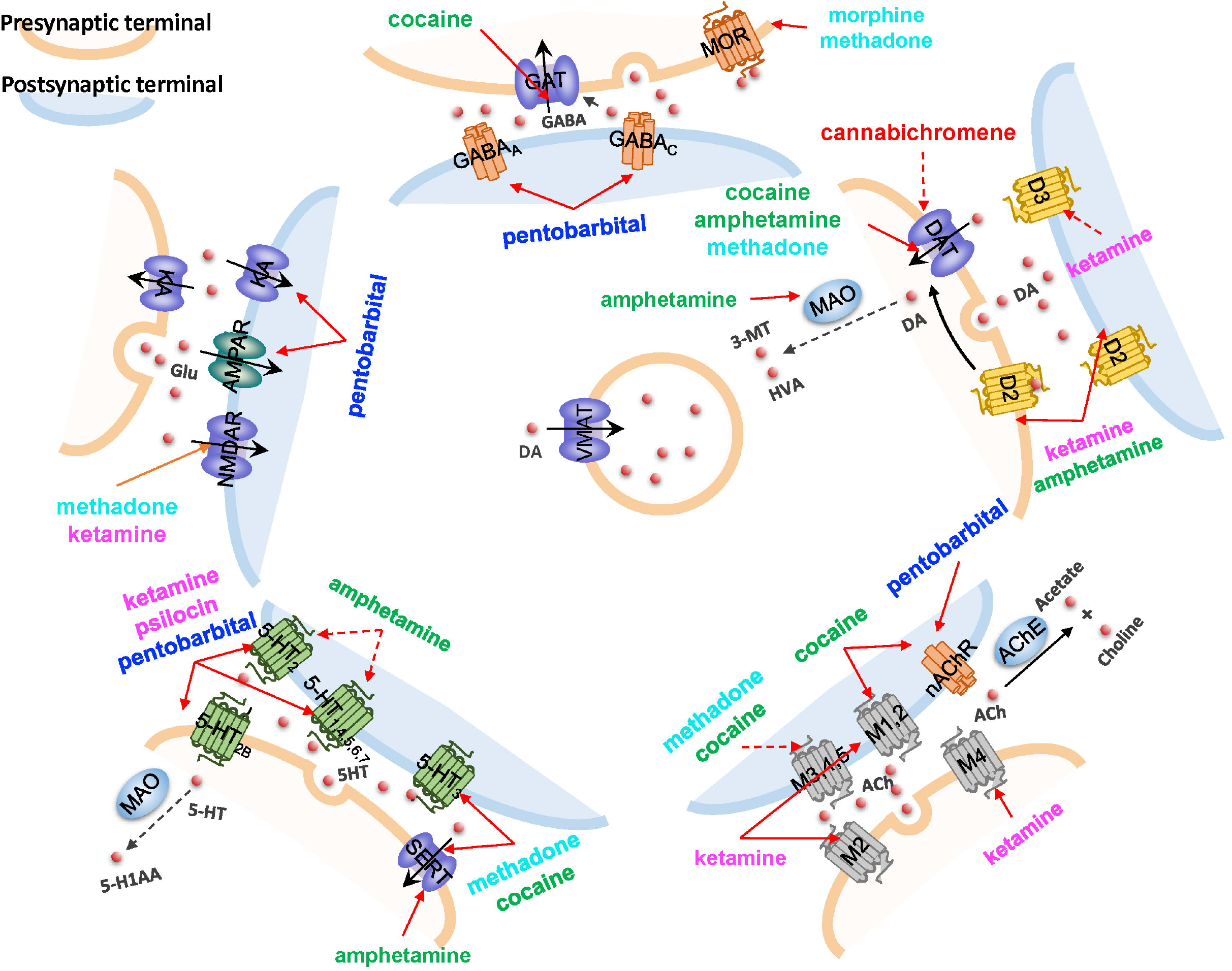
The impact of drugs of abuse on synaptic neurotransmission. Five major neurotransmission events are highlighted, mediated by (*counterclockwise, starting from top):* GABA receptors and transporters, ionotropic glutamate receptors (NMDAR and AMPAR) and cation channels, serotonin (5HT) receptors (5-HTR) and transporters (SERT), muscarinic or nicotinic AChRs, and dopamine (DA) receptors and transporters. Vesicular monoamine transporters (VMAT) that translocate DA are also shown. Drugs affecting the different pathways are listed, color coded with their categories, as presented in **Figure 1**. *Solid red arrows* indicate a known drug-target interaction, *dashed red arrows* indicate predicted drug-target interactions. Other molecules shown in the diagram are: KA, kainate receptor; MAO, monoamine oxidase; HVA, homovanillate; 3-MT, 3-methoxytyramine; MOR, mu-type opioid receptor; AChE, acetylcholinesterase; and 5-H1AA, 5-hydroxyindoleacetate.

It is worth noting that predictions by our PMF model lead to a better understanding of the way drugs of abuse affect neurotransmissions. In addition to the new role of M5 we discussed in Section 3.2, our PMF model predicted that cannabichromene, a cannabinoid whose primary target is the transient receptor (TRPA1), is found to interact with DAT and directly regulate dopaminergic transmission, which will require further examination. Synaptic neurotransmission events act as upstream signaling modules that ‘sense’ the early effects of drug abuse. In the next section, we focus on the downstream signaling events elicited by drug abuse.

### 3.6 mTORC1 emerges as a downstream effector activated by drugs abuse

The calcium-, cAMP-, Rap1-, Ras-, AMPK-, ErbB-, MAPK-, and PI3K-Akt-signaling pathways in the SG category (**Supplementary Table 4**) crosstalk with each other and form a unified signaling network. As shown in **Figure 5**, ligand-binding to GPCRs modulates the production of cAMP, which leads to the activation of Rap1. Activated Rap1 modules the Ca^2+^ signaling by inducing the production of inositol triphosphate (IP_3_) and also activates the PI3K-Akt signaling cascade. Stimulations of ErbB family of receptor tyrosine kinases (related to epidermal growth factor receptor EGFR) as well as insulin-like growth factor receptor IGF1R trigger both PI3K-Akt and MAPK signaling cascades (proteins colored *blue* in **Figure 5**). Notably all these pathways merge and regulate a group of downstream proteins (shown in *dark yellow* in **Figure 5**); and at the center of this cluster lies the mammalian target of rapamycin (mTOR) complex 1 (mTORC1) which is likely to be synergistically regulated by all these merging pathways.

**Figure 5.**
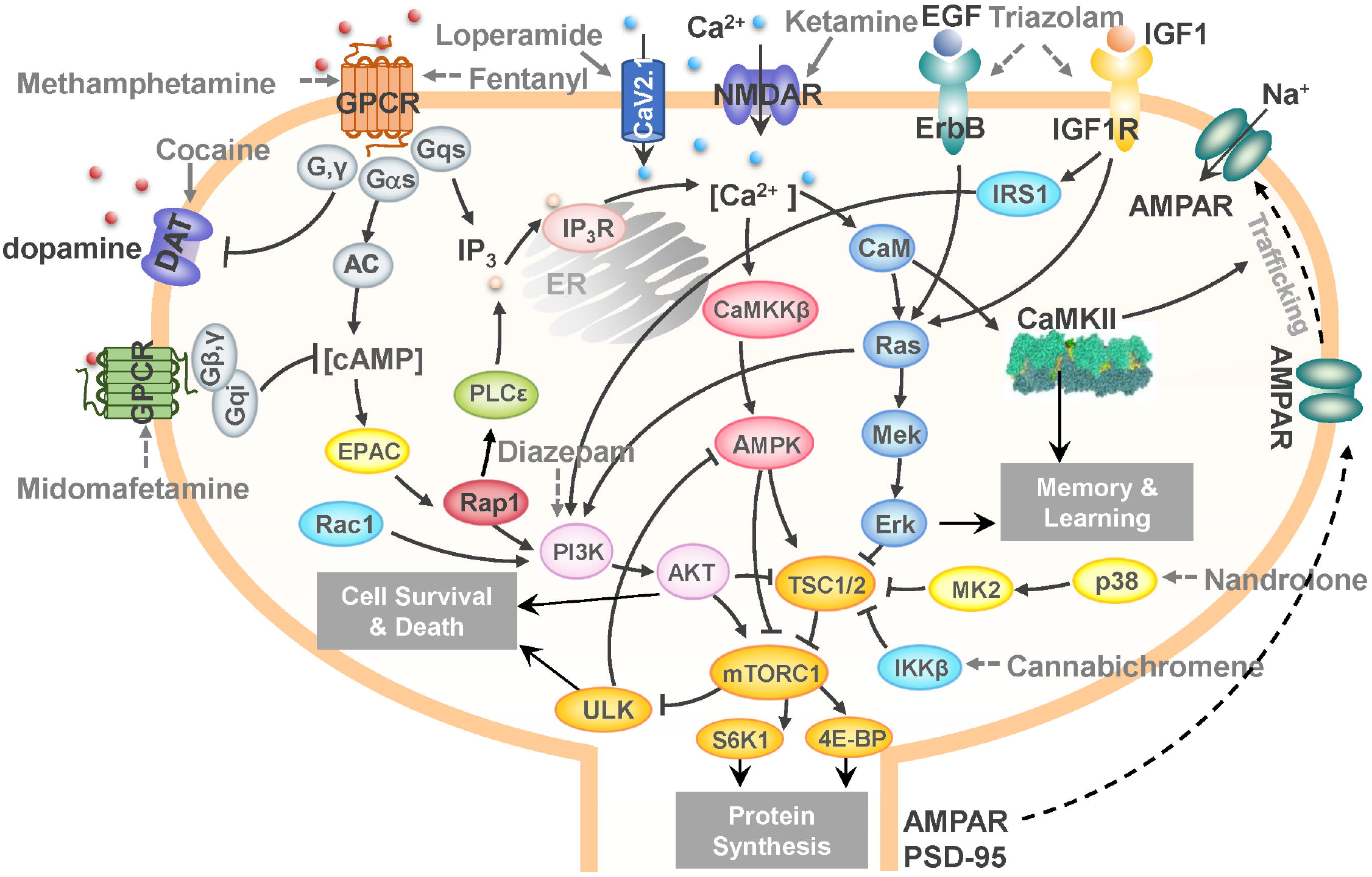
A unified signaling network mediates the effects of drugs of abuse. *Black arrows* represent the activation, inhibition, and translocation events during signal transduction. *Solid gray arrows* represent the known drug-target interactions. *Dashed gray arrows* represent predicted drug-target interactions. The diagram illustrates the targets of several drugs of abuse belong to different categories: loperamide and fentanyl belong to the opioids group; midomafetamine and ketamine are from the hallucinogens group; triazolam and diazepam are CNS depressants; cannabichromene is a cannabinoid; methamphetamine and cocaine are CNS stimulants; nandrolone is an anabolic steroid. The mTORC1 emerges as a hub where the effects on several targets of abused drugs appear to be consolidated to lead to cell death and/or protein synthesis in the CNS, and in particular AMPAR/PSD95 synthesis that induces morphological changes in the dendrites.

mTORC1 is not only a master regulator of autophagy (Rabanal-Ruiz et al., 2017), but also controls protein synthesis and transcription (Ma and Blenis, 2009). It has been reported to promote neuroadaptation following exposure to drugs of abuse including cocaine, alcohol, morphine and Δ^9^-tetrahydrocannabinol (THC) (Neasta et al., 2014). Our results suggest that mTORC1 may act as a universal effector of the cellular response to drug abuse at an advanced (preoccupation and anticipation, or craving) stage, controlling the synthesis of selected proteins and ensuing cell growth, which may result in persistent alterations in the dendritic morphology and neuronal circuitry.

In **Figure 5**, selected interactions between drugs from different substance groups and their targets are highlighted using *gray* arrows. Our PMF model predicted that diazepam would interact with PI3K to influence mTORC1 signaling (*dashed gray arrows* denote predictions). It has been reported that Ro5-4864, a benzodiazepine derivative of diazepam suppresses activation of PI3K (Yousefi et al., 2013), which corroborates our prediction. We further predicted that cannabichromene may interact with IκB kinase ß (IKKß) to regulate mTORC1 by inhibiting TSC1/2. Interestingly, another cannabinoid arachidonoylethanolamine is known to directly inhibits IKKβ (Sancho et al., 2003). Taken together, our results identified a unified network that underlies the development of drugs addiction, in which mTORC1 appears to play a key effector role.

## 4 Discussion

In the present study we focused on the targets and pathways affected by drugs of abuse, toward gaining a systems-level understanding of key players and dominant interactions that control the response to drug abuse and the development of drug addiction. Using machine learning methods, we focused on 50 drugs of abuse that form a chemically and functionally diverse set, and analyzed their 142 targets as well as the corresponding cellular pathways and their crosstalk. Our analysis identified:

i. 48 additional proteins targeted by drugs of abuse, including PIK3CA, IKBKB, EGFR, and IGF1R, are shown to be key mediators of downstream effects of drug abuse.
ii. 161 new interactions between the drugs of abuse and the known and predicted targets, including those between cocaine and M5, methylphenidate and OPRM1, and diazepam and PI3K, not reported in existing DBs, but supported by prior experiments, and others (e.g. the interactions of canabichromene with IKBKB and DAT) that await experimental validation.
iii. A dataset of 70 pathways, composed of 6 neurotransmission pathways, 46 signal transduction pathways, 8 neuroplasticity pathways and 10 autonomic nervous system innervation pathways which are proposed to govern different stages of the molecular, cellular and tissue level responses to drug abuse and in addiction development.

Overall, our comprehensive analysis led to new hypotheses on drug-target interactions and signaling and regulation mechanism elicited by drugs of abuse in general, along with those on selected targets and pathways for specific drugs. Below we elaborate on the biological and biomedical implications of these findings.

### 4.1 Persistent restructuring in neuronal systems as a feature underlying drug addiction

Enriched pathways in the neuroplasticity category include gap junction, LTP, LDP, adherens junction, regulation of actin cytoskeleton, focal adhesion, axon guidance, and tight junction (**Supplementary Table 4**). These are responsible for the changes in the morphology of dendrites. For instance, DA regulates excitatory synaptic plasticity by modulating the strength and size of synapses through LTP and LTD (De Roo et al., 2008; Volkow and Morales, 2015). The restructuring of dendritic spines involves the rearrangements of cytoskeleton and actin-myosin (Volkow and Morales, 2015). The axon guidance molecules guide the direction of neuronal growth.

Drugs of abuse can induce the changes in CNS through these pathways. For example, chronic exposure to cocaine increases dendritic spine density in medium spiny neurons (Russo et al., 2010). The disruption in axon guidance pathway and alteration in synaptic geometry can result in drug-related plasticity (Bahi and Dreyer, 2005). The persistent restructuring in the CNS caused by drugs of abuse is responsible for long-term behavioral plasticity driving addiction (Volkow et al., 2003; Russo et al., 2010; Volkow and Morales, 2015). As will be further discussed below, mTORC1 plays a central role in the synthesis of new proteins (e.g. AMPARs) and thereby neuronal (dendrites) growth, alteration of the synaptic geometry and therefore rewiring of the neuronal circuitry.

### 4.2 ANS may mediate the negative-reinforcement of drug addiction

Our results further show that the pathways regulating ANS-innervated systems are associated with drugs of abuse. As the NP pathways may influence the neuroplasticity in the ANS, we hypothesize that drugs of abuse may induce the persistent restructuring in ANS as well. The drug-related plasticity in ANS may lead to the dysregulation of ANS-innervated systems and cause negative effects and feelings during the second stage of drug addiction.

Drug addiction is well known as a brain disease (Volkow and Morales, 2015). However, many drugs of abuse can disrupt the activity of ANS and cause disorders in ANS-innervated systems (Al-Hasani and Bruchas, 2011; Huang, 2017). For example, opioids (e.g. morphine) alter neuronal excitability and neurotransmission in the ANS (Wood and Galligan, 2004), and induce disorders in gastrointestinal system, smooth muscle, skin, cardiovascular, and immune system (Al-Hasani and Bruchas, 2011). Cannabinoids (e.g. THC) modulate the exocytotic NE release in ANS-innervated organs through presynaptic cannabinoid receptors (Ishac et al., 1996).

The pathways we identified in the ANS category regulate insulin secretion, gastric acid secretion, vascular smooth muscle contraction, pancreatic secretion, salivary secretion and renin secretion (**Supplementary Table 4**). Their dysfunction may be associated with the autonomic withdrawal syndrome, such as thermoregulatory disorder (chills and sweats) and gastrointestinal upset (abdominal cramps and diarrhea), which has been observed in drug/substance users (Wise and Koob, 2014). In addition, the stress and depression caused by these negative effects may be part of the negative reinforcement of drug addiction (Self and Nestler, 1995; Koob and Le Moal, 2001). In other words, the drug induced ANS disorders can feedback to CNS and mediate the negative reinforcement. Compared to the structural changes in CNS, the disorder and persistent restructuring in ANS is less studied and it could be a future direction in the study of development of drug addiction and related diseases.

### 4.3 mTORC1 signaling plays a key role in mediating cellular morphological changes in response to continued drug abuse

The functioning and regulation of mTOR signaling has been elucidated over the past two decades. It became clear that mTORC1 plays a crucial role in regulating diverse cellular processes including protein synthesis, autophagy, lipid metabolism, and mitochondrial biogenesis (Saxton and Sabatini, 2017). In the brain, mTORC1 coordinates neural development, circuit formation, synaptic plasticity, and long-term memory (Lipton and Sahin, 2014). The dysregulation of mTORC1 pathway is associated with many neurodevelopmental and neurodegenerative diseases such as Parkinson’s disease and Alzheimer’s disease. mTORC1 been noted to be an important mediator of the development of drug addiction and relapse vulnerability (Dayas et al., 2012). Accumulating evidences show that pharmacological inhibition of mTORC1 (often through rapamycin treatment) can prevent sensitization of methamphetamine-induced place preference (Narita et al., 2005), reduce craving in heroin addicts (Shi et al., 2009), attenuate the expression of alcohol-induced locomotor sensitization (Neasta et al., 2010), suppress the expression of cocaine-induced place preference (Bailey et al., 2012), protect against the expression of drug-seeking and relapse by reducing AMPAR (GluA1) and CaMKII levels (James et al., 2014), and inhibit reconsolidation of morphine-associated memories (Lin et al., 2014).

Our unbiased computational analysis based on a diverse set of 50 drugs of abuse supports the hypothesis that mTORC1 may act as a universal effector or controller for neuroadaptations induced by drugs of abuse (Neasta et al., 2014). The major signal transduction pathways we identified that involve targets of drugs of abuse interconnect and converge to the mTORC1 signaling cascade (**Figure 5**). Most drugs of abuse in our list target upstream regulators of mTORC1, including membrane receptors (e.g. GPCRs, RTKs and NMDAR), kinases (e.g. PI3K, p38α, and IKKβ), and ion channels (e.g. Ca_V_2.1 and TRPV2). Notably, the drug-related impact of some of these targets has been experimentally confirmed. For example, blockade of NMDAR using MK801 reduces the amnesic-like effects of cannabinoid THC (Puighermanal et al., 2009). Likewise, PI3K inhibitor LY294002 can suppress morphine place preference (Cui et al., 2010) and the expression of cocaine-sensitization (Izzo et al., 2002).

The downstream effectors of mTORC1, which specifically mediate drug behavioral plasticity is far from known. mTORC1 can mediate the activation of S6Ks and 4E-BPs, which leads to increased production of proteins required for synaptic plasticity including AMPAR and PSD-95 (Dayas et al., 2012). EM reconstruction of hippocampal neuropil showed the variability in the size and shape of dendrites depending on synaptic activity (Bartol Jr et al., 2015), which in turn correlates with information storage. Recently studies have revealed that Atg5- and Atg7-dependent autophagy in dopaminergic neurons regulates cellular and behavioral responses to morphine (Su et al., 2017). Cocaine exposure results in ER stress-induced and mTORC1-dependent autophagy (Guo et al., 2015). Fentanyl induces autophagy via activation of ROS/MAPK pathway (Yao et al., 2016). Methamphetamine induces autophagy through the ĸ-opioid receptor (Ma et al., 2014). These observations are all consistent with the conclusion drawn here with regard to the role of mTORC1 as a major effector of cellular responses to drug addiction.

### 4.4 Drug repurposing opportunities for combatting drug addiction

Autophagy modulating drugs have been shown to have therapeutic effects against liver and lung diseases. The signaling network presented in **Figure 5** involves many targets of such drugs. For instance, carbamazepine affects IP_3_ production and enhances autophagy via calcium-AMPK-mTORC1 pathway (Hidvegi et al., 2010). It has been identified as a potential drug for treating α1-antitrypsin deficiency, hepatic fibrosis, and lung proteinopathy (Hidvegi et al., 2010; Hidvegi et al., 2015). Rapamycin is a potential drug for lung disease such as fibrosis (Abdulrahman et al., 2011; Patel et al., 2012). Other liver and lung drugs which facilitate the removal of aggregates by promoting autophagy may also affect drug-related neurodegenerative disorders. **Supplementary Table 7** summarizes 15 autophagy-modulating drugs for liver and lung diseases. Target identification and pathway analysis of this subset of drugs using the same protocol as those adopted for the 50 drugs of abuse indeed confirmed that drugs of abuse and liver/lung drugs share many common pathways (**Supplementary Figure 5**). Notably, among those pathways, neuroactive ligand-receptor interactions, calcium signaling, and serotonergic synapse pathways are among the top 10 enriched pathways of both drugs of abuse and liver/lung drugs. Amphetamine addiction and alcoholism are also enriched by targets of liver/lung drugs. Thus, an interesting future direction is to examine whether autophagy modulating drugs for liver and lung diseases could be repurposed, if necessary by suitable refinements to increase their selectivity, for treating drug addiction.

## 6 Author Contributions

FP, HL and IB conceived and designed the research. FP and HL performed the research. FP, HL, BL and IB analysed the results. FP, HL, BL and IB wrote the manuscript.

## 8 Acknowledgments

FP wish to thank Dr. D. Lansing Taylor for his mentorship.

